# Developmental Continuity of Brain Network Core Organization in *C. elegans*

**DOI:** 10.64898/2026.06.12.730308

**Authors:** Prateek Yadav, Aradhana Singh

## Abstract

The brain is the most captivating chef d’oeuvre of nature. Naturally then, the mind wonders about the process that births such a fascinating organ. Neurodevelopment is a complex yet robust phenomenon that conceals answers to our questions in its intricacies. In an attempt to shed some light on this matter, we study the developing brain connectome of the nematode, *C. elegans* across the post-embryonic phase. A tiny organism with only around 200 neurons comprising its brain and yet a diverse array of behaviors to display, it makes for a great model. Starting with most of its head neurons already present at hatching, the worm brain accumulates numerous more synaptic connections increasing the edge density. It maintains a weak connectivity throughout thereby, balancing global communication as well as hierarchy. At the mesoscopic level, we find that the core has a conserved backbone of persistent neurons along with a dynamic component formed of transient/recurring neurons. Moreover, the connectome has a rich club organization since the early stage which selectively strengthens indicating progressively denser connectivity among the integrators due to the previously reported asymmetric synapse addition. This asymmetry also shows up in the preservation of input hubs across development and the progressively more centralized organization of the in-degree k-core. Our work provides a new perspective into the neurodevelopment of the brain that may facilitate our understanding of its functioning.

## 1 Introduction

The study of brain development can offer insights into the function of the brain and how and why neurodevelopmental disorders appear [1]. This would lead us neuroscientists closer to the north star of the field of network neuroscience that is, finding the link between structure and function [2]. One way this can be approached is by studying the mesoscopic organization of the developing network because that allows us to study the emergent properties of the complex network that the brain is [3, 4]. Comparing patterns of neurodevelopment across taxa can be informative as to how generalized the process is and what purpose the unique attributes serve any organism [1].

In terms of post-embryonic neural remodeling, the seminal work on *C. elegans* biology by Sulston and his group had reported that certain dopaminergic neurons are generated after hatching [5]. Since then, we have developed a more comprehensive understanding of post-embryonic development through this highly tractable model organism. This includes an exceptional case of transdifferentiation wherein epithelial cells transform into neurons [6, 7] tail glia cells transmute into male-specific neurons[8]. Besides transdifferentiation, certain glial cells have also been observed to produce neurons in the post-embryonic phase, for example, in males, AMso divides to form neurons [9]. The transformation of larva into dormant dauer also involves arborization of some sensory neurons [10–12]. The neural network formed of the gap junctions is known to be adaptive toward environmental fluctuations [13]. Significant remodeling has also been observed in the circuits involving motor neurons [14, 15]. Furthermore, sex-specific rewiring also occurs in the worm, for instance, the synapse elimination between PHB and AVG neurons in hermaphrodites [15]. Interestingly, it has been suggested that the notable level of neuronal plasticity actually helps to establish stereotypic connectivity [16].

These postnatal cellular changes are closely associated with corresponding shifts in behavior. While certain activities like chemotaxis and locomotion [17, 18] are fine-tuned as the worm matures into an adult, there’s also the adoption of several novel behaviors like egg-laying, mate searching, mating-specific locomotion, and pheromone-mediated social interactions [19–22]. Controllability analysis has revealed that as *C. elegans* develops, its brain network becomes increasingly controllable at the global level, whereas control over muscle-specific outputs diminishes, suggesting a reallocation of regulatory capacity from fundamental locomotion to more complex behaviors and decision-making processes [23].

*Caenorhabditis elegans* is a common 1mm long nematode found in the soil that feeds on decaying plant matter [24, 25]. It has only 302 somatic neurons and yet displays many distinct behaviors, thus making it a great organism to explore the structure-function paradigm [26]. Technological advancement has made it possible to reconstruct the connectome of model organisms like the worm and the fruit fly. In the case of the worm, we have come far from the pioneering work by White et al. in 1986 [26]. Today, we have reconstructions of the entire somatic nervous system of both sexes of *C. elegans* [27] in addition to the pharyngeal connectome of the hermaphrodite worm [28]. The brain of the worm across its development has been mapped as well, forming the basis of this study [29]. Even the chemical connectome of the nerve ring, the major part of the worm brain, of the dormant stage, dauer, was reconstructed [30]. Beyond the chemical and electrical synapses, the network of extrasynaptic signaling pervasive in the complete nervous system of the worm has also been delineated [31]. Furthermore, there is a great information available regarding the biology of the organism, which includes gene expression profiles, lineage, etc

In 2021, Witvliet et al. mapped the connectome of *C. elegans* at 7 different timepoints in its development revealing certain principles of brain development. They found that the modularity increases with age and the brain network becomes more feedforward with maturation. Interestingly, they observed that synapse addition is not uniform in that more synapses are added to existing input connections of hub neurons, and more synapses appear to establish new input connections for hub neurons. Moreover, from the perspective of a cell, in general, the input connections strengthen in an uncorrelated manner across development, whereas its output connections strengthen in proportion to each other. We wanted to study the upshot of this at the mesoscale and macroscale levels of the network. Another study on this same dataset and found that the degree distribution of the worm brain decays exponentially, and consistently so, throughout its post-embryonic development [32]. This means that the probability of finding high-degree hubs is less than a scale-free regime.

Our study examines the mesoscopic architecture of the worm brain throughout its post-embryonic development by employing core decomposition and rich club analyses with the aim of uncovering insights into its neurodevelopment. We study the connectome at 7 different timepoints in its development and study the parametric trends. The network gradually becomes denser and is always wired in a way to promote weak connectivity, balancing global communication and hierarchy. Examining the core layers using a comprehensive approach, we observe that the connectome has a group of neurons that invariably are part of the core throughout development when each core type is considered separately. Additionally, with development, the core composition shifts to accommodate different cells (embryonic and post-embryonic), which may be involved in a new function or account for behaviors unique to that individual. This suggests that the developing brain with a stereotypical character also allows for a dynamic core. This complex core organization led us to explore whether core participation is influenced by lineage proximity among the neurons; we find that it plays a minor role and thus, postulate that there are other factors playing a role in tandem with lineage proximity to drive core formation. Moreover, we find that since early on in its development, the brain is wired to balance global communication and integration of information. This quality as a whole sustains till maturation; however, by a weighted-in-degree metric, the effect increases with development, likely due to asymmetric synapse addition. Notably, the input hubs are preserved across development, and the topology around the in-degree k-core progressively becomes more centralized. Thus, we extend the findings of Witvliet et al. [29] by showing that a directional asymmetry appears at the meso- and macro-scale.

## 2 Results

### 2.1 Weakly Connected Brain Connectome achieves high Connection Density with Development

There are 8 networks representing 7 different stages in the development of the worm’s brain. The nerve ring neuropil, along with the ventral ganglion, is considered to be the brain of the worm. The adult stage is mapped twice. Each network is directed and weighted by the number of chemical synapses. We probe the developing brain network by primarily using core decomposition analyses, rich club analysis, and modular data in addition to other supporting network science tools. The brain network is weakly connected throughout development and never strongly connected (Table 1). A directed network is weakly connected if there’s a path from one node to every other node, disregarding the directionality of the edges, while it’s a strongly connected network if every node is reachable from every other node by following the directed edges. Real-world networks are known to lack strong connectivity because of their hierarchical structure, since it’s the backward edges that are necessary to attain a strong connectivity [33, 34]. By maintaining weak connectivity, the brain assures global communication without compromising on hierarchy.

**Table 1:** Network properties of Worm Brain at different stages. *N*_*c*_, *N*_*n*_, *E, S, ρ, ρ*_*n*_, *k*, WC, SC, *w* represent the number of all cells, the number of neurons, the number of edges, number of synapses (chemical and neuromuscular junctions), connection density of the entire network, connection density of the neuronal subnetwork, state of being weakly connected, state of being strongly connected, and average connection weight.

| Stage | $N_c$ | $N_n$ | $E$ | $S$ | $\rho$ | $\rho_n$ | $\langle k \rangle$ | WC | SC | $\langle w \rangle$ |
| --- | --- | --- | --- | --- | --- | --- | --- | --- | --- | --- |
| L1<sub>0h</sub> | 209 | 167 | 775 | 1296 | 0.0178 | 0.02435 | 7.42 | True | False | 1.672 |
| L1<sub>5h</sub> | 209 | 167 | 986 | 1895 | 0.0227 | 0.0312 | 9.44 | True | False | 1.922 |
| L1<sub>8h</sub> | 210 | 168 | 1012 | 2128 | 0.0231 | 0.03162 | 9.64 | True | False | 2.103 |
| L1<sub>16h</sub> | 219 | 177 | 1136 | 2777 | 0.0238 | 0.03245 | 10.37 | True | False | 2.445 |
| L2 | 222 | 180 | 1515 | 4116 | 0.0309 | 0.04109 | 13.65 | True | False | 2.717 |
| L3 | 223 | 181 | 1525 | 4456 | 0.0308 | 0.04036 | 13.68 | True | False | 2.922 |
| Ad<sup>1</sup> | 223 | 181 | 2191 | 7455 | 0.0443 | 0.05933 | 19.65 | True | False | 3.403 |
| Ad<sup>2</sup> | 223 | 181 | 2186 | 7970 | 0.0442 | 0.05933 | 19.61 | True | False | 3.646 |

With maturation, the edge density of the brain progressively increases(Table 1). The worm undergoes continuous development [35] and the brain growth, as measured by total neurite length, scales proportionally with overall body length [29]. The synapse number increases in proportion to these as well [29]. While the number of neurons marginally increases with maturation because, although the worm undergoes two bursts of neurogenesis, most of the neurons in the head neuropil are born in the first phase, i.e., before hatching [36]. Subsequently, the new synapses predominantly strengthen the existing connections as attested by the increase in average connection weight (Table 1). Moreover, the high density of connections enables the multifunctional neurons [37] to take part in multiple circuits to execute diverse behaviors. Furthermore, we find that strong connectivity increases with development, as evidenced by the fraction of the size of the giant strongly connected component. This is surprising because Witvliet et al. found that the network becomes more feedforward, so it would be expected that strong connectivity decreases [38].

### 2.2 Core Decomposition

#### 2.2.1 The Brain Network displays a Consistent Core Composition

The k-core decomposition reveals that both in-degree and out-degree k-core numbers remain relatively small (Figure 1 (B)). However, larger k-core values are observed in later developmental stages, which align with an increasing number of connections (node degrees) as the connectome matures. The stacked bar plots for cell type counts indicate that diverse neuron types contribute to these cores, reflecting structural stability and consistent connectivity patterns across neuron types. In contrast, the s-core decomposition highlights that core numbers, based on strength, are significantly higher than those observed in the k-core decomposition at all stages. The disparity between k-core and s-core numbers is indeed related to how new synapses are added during development. Synapse addition prefers to selectively strengthen a cell’s individual connections rather than creating entirely new ones, as evidenced by the steady increase in average connection weight (Table 1).

**Figure 1:**
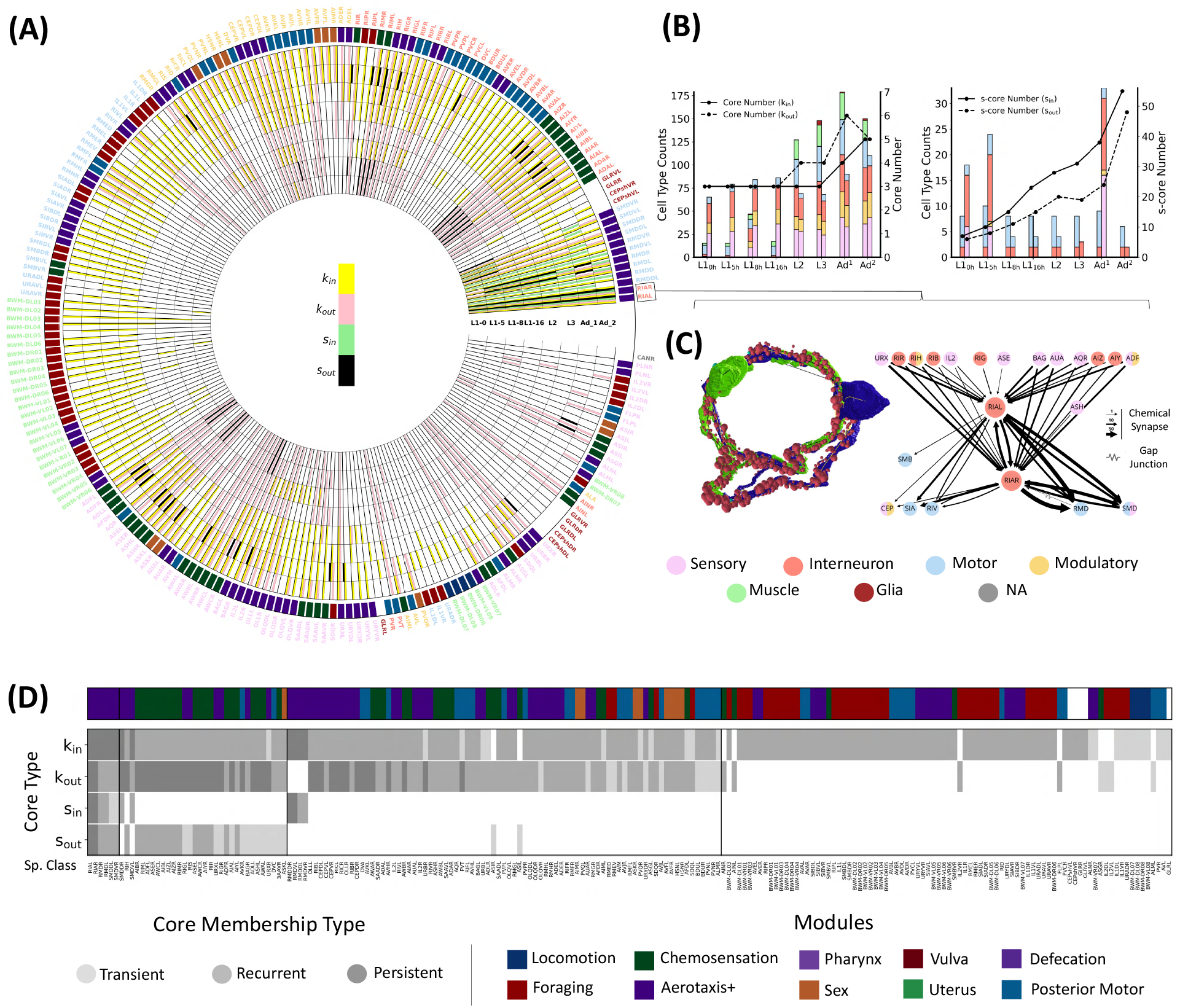
k-core and s-core decomposition of the *C. elegans* brain networks across development. (A) Radial colormap showing core participation of *C. elegans* cells across stages and core decomposition methods. Each concentric ring represents a developmental stage (from inner to outer: *L*1_0*h*_ to *Ad*^2^). Each radial segment corresponds to a cell-specific class label colored by cell type. Colored outer ticks indicate behavioral modules defined in Emmons 2024 [37]. The presence of a neuron in the core at a given stage is shown by a filled radial cell subdivided into four angular wedges. Each wedge indicates membership in one of the four core types. (B) Stacked bar plots showing the distribution of cell types within the in-degree and out-degree cores (k-core-left panel; s-core-right panel) across stages. For every stage, the left bar corresponds to the in-degree core and the right bar to that of the out-degree core. The overlaid line represents the main core number. (C) Left Panel: Anatomical representation of RIAL (green) and RIAR (blue) neurons with synapses shown in red. Right Panel: the subgraph of RIA neurons (Courtesy NemaNode [29]). Color code for the cell types in (A), (B), and (C) is provided. (D) plots all the cells that ever feature in the core during the course of development into 3 categories: transient, recurrent, and persistent for each of the four core types. The cells are ordered in a descending manner according to the number of core types they are ever part of. Color code for modules shown in (A) and (D) is provided.

This selective strengthening results in fewer new connections per node, which keeps the degree-based k-core numbers relatively lower. At the same time, the cumulative increase in synaptic weights leads to significantly higher s-core numbers. Thus, while the k-core reflects the modest addition of new connections, the s-core captures the cumulative strengthening of existing connections, accounting for the observed differences.

Heterogeneity in terms of connection strength facilitates efficient computation [39, 40]. Thus, it is important to examine strong connections, and s-core decomposition is a good tool for the same. S-core decomposition unveils circuits involving RMD, SMD, and RIA neurons (Figure 1 (A)). These all belong to the community of Aerotaxis (and other unassigned functions). Individually, SMD and RMD are both involved in locomotion. RMD neurons are specifically known to play a role in spontaneous foraging movements and the head withdrawal reflex [41]. RIA (Figure 1 (C)) are second-layer interneurons known to be part of the thermotaxis core and involved in local search and dispersal movements [42]. The two RIA neurons are found to be members of all types of cores, which is not surprising given that *C. elegans* is known to display bilateral symmetric pairing homophily [43].

To study core participation across development, for each core type, we categorized the participating cells as transient, recurrent, or persistent if they appear in the core once, a few times, or all the time across development, respectively (Figure 1 (D)). Firstly, we see that a few cells involved in aerotaxis (and other unassigned functions) tend to appear in all 4 core types. An older study on the binarized, undirected network of the somatic nervous system of the worm had found that all hubs are born by 450 mins after fertilization (before first obvious signs of any motor activity) [44]. Narrowing down to the head neurons, we find that only AIBR, RIBL, and RIAR are persistent in at least one core type in the brain network alone. Notably, similar to the above-mentioned study, we observe that all the 51 cells that are persistent for at least one core type are born early, before any twitching is observed. However, this is not surprising in the context of the brain since the vast majority of its cells are born before that time point. However, most of the core neurons (for any core type) belong to the bracket of transient/recurrent. A few among these are simply born late and either immediately or eventually form a part of the core; for example, the AVF neurons are born around 1675 minutes after fertilization. They are involved in sexual circuits of both the male and the hermaphrodite, so it could be interesting to see if they emerge as a hub even in the male connectome [27, 37]. AVF neurons are noted for their role in the momentary burst of forward movement before egg-laying by the hermaphrodite worm [27, 37]. But many of them are born early; however, they are only later integrated into the brain network and eventually the core. For instance, HSN neurons undergo a slow maturation process. They are born in the tail region around 410-420 minutes after fertilization, they later migrate to the midbody by the L1 stage, initiate axon outgrowth in the L3 stage, and form synapses in L4, and finally become a functional adult brain network [45, 46]. Some of these neurons could also be pointing to variation in behavior across stages, as it has been found that *C elegans* olfactory behavior varies with stage [47]. Adult *C elegans* have been found to strongly respond to diacetyl, a food odorant, while the larvae are weakly attracted to it. This change is brought about in germline cells [47]. Perhaps many other transient or recurrent neurons belonging to either of the k-cores are involved in foraging during later stages of the development, as there have been changes observed in foraging behavior too in the worm [48].

However, it is also likely that many of these transient or recurrent neurons are simply artifacts of the within-individual differences in neuronal architecture. It’s known that roughly 16% of the chemical synapses are not conserved even among isogenic individuals [29]. The strongest evidence for this could be the cyclical appearance-disappearance of cells in the core because the literature doesn’t support any reason as to why a certain cell would behave as a hub at one point, get demoted, and then emerge as a hub again. Particularly, the lack of similarity in the arrangement of core shells between the two adults could be due to the differences in behavior that are known to arise even in isogenic animals reared in the same environment [49–51].

All the neurons forming the s-cores at any stage are embryonic, possibly because s-cores are composed of those cells with high weighted degrees and thus, early born neurons have the most time to gain from the synaptic strengthening, which is also biased toward hubs [29].

#### 2.2.2 No evidence for a multi-core structure in the Worm Brain

To investigate whether core decomposition methods provide a sufficient outlook on the cohort of hubs, we implemented checks for the presence of multicores, that is, other cohorts of hubs that might be languishing in lower shells and thus, left undiscovered by conventional core decomposition techniques. A heatmap showcases the overlap of core-adjacent shells corresponding to different core decomposition methods with different modules in the network (Figure 2). There’s no pattern consistent across development for any core type, although there are a few scattered instances of certain core-adjacent shells showing strong enough overlap with any one of the modules for them to be considered subsidiary cores. However, due to the lack of any emerging trend across development, we consider the cores obtained through core decomposition methods to be adequate for studying hub populations.

**Figure 2:**
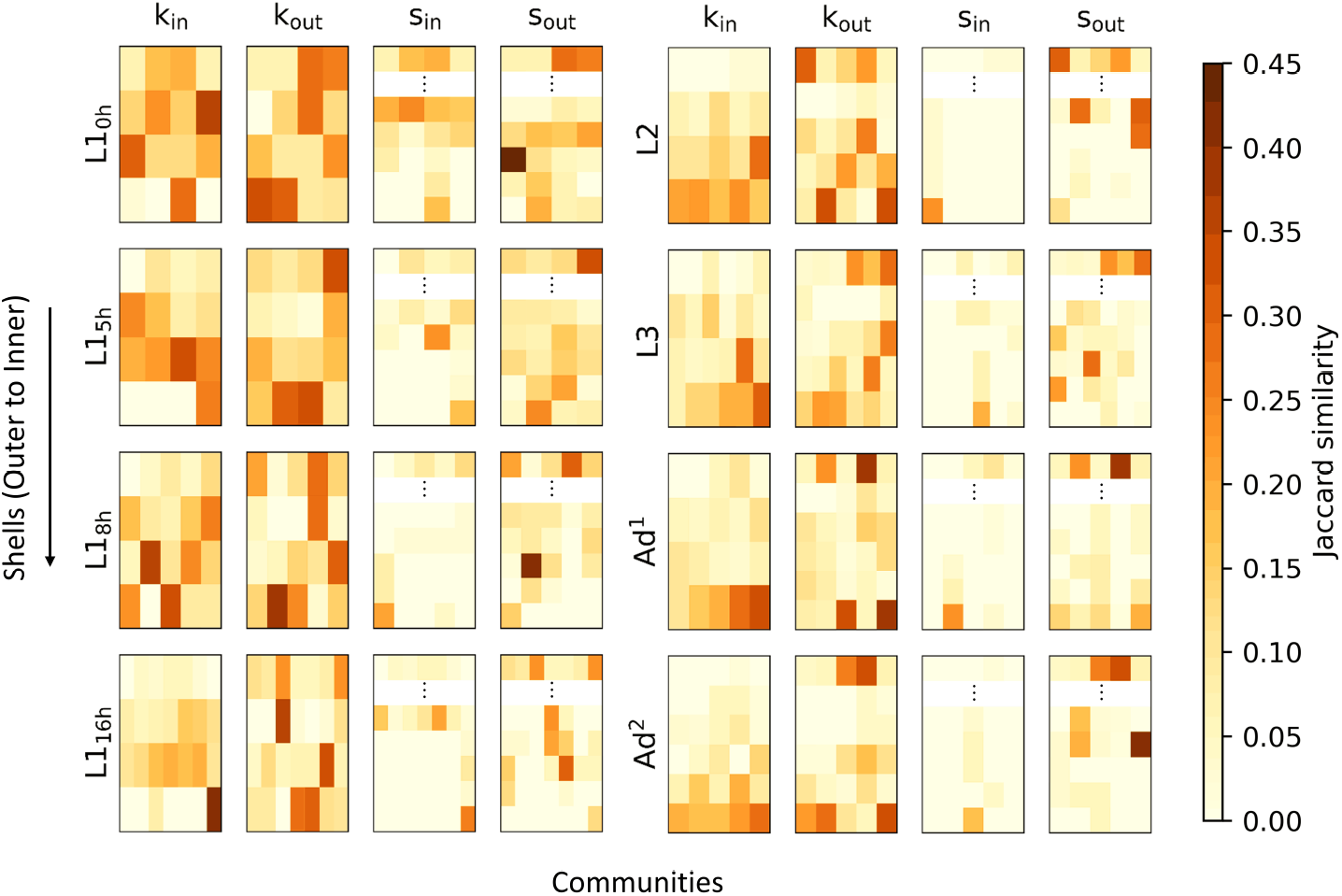
Investigation of a multicore structure corresponding to each core decomposition method. Each cell in a heatmap shows the Jaccard similarity between the corresponding shell and the corresponding module. For ease of visualization, the outer s-cores are not shown and represented by a vertical ellipsis.

#### 2.2.3 Lineage Proximity: Core Cells are Weakly Developmentally Clustered

To see how developmentally clustered the core participation of neurons is, we studied the lineage distance distribution of the core neurons. It’s been known that lineage-based homophily positively influences synaptic connectivity, likely by increasing the chance of spatiotemporal proximity [43]. We replicated the analyses for the k-cores of the brain network at all stages. Our results suggest that during the early stages, the influence of lineage on the connectivity, if there’s any at all, is not statistically detectable, perhaps due to noise (Figure 3). On the other hand, the adult stages show a moderately lineage-dependent synaptic connectivity for both in- and out-degree k-cores. Intrigued, we wish to see the pattern for the overall network connectivity (unweighted). We find that it seems to be progressively becoming more negatively influenced by the lineage distance, suggesting that the likelihood of two closely related neurons synapsing together increases with development (Figure 3). Interestingly, the connectivity seems to be more permissive in terms of lineage-related constraints in the early larval stages, given the weaker association. This indicates that perhaps, in the synapse formation in the embryonic stage is more influenced by factors other than lineage proximity. One factor that could be responsible is the neuronal activity, which has been found to promote synapse formation [52, 53]. Additionally, it’s crucial to understand that, like previous work on mouse neocortex shows, lineage selectively plays a role in neuronal connectivity: clonally related neurons have a greater likelihood to wire together vertically across cortical layers; however, their lineage proximity doesn’t influence the chances of establishment of connections within the same layer. This suggests that functional circuits arise from a combination of clonal and nonclonal connectivity [54]. Moreover, it’s known that neuronal hubs have a correlated transcriptional signature, which is mediated neither by lineage proximity nor by any factor among birth time, cell type, and distance, etc. [55]. Rather, it could be inherently related to their respective identities. On top of all this, our results offer ideas for future research into how different factors vary their influence on synaptic connectivity throughout development.

**Figure 3:**
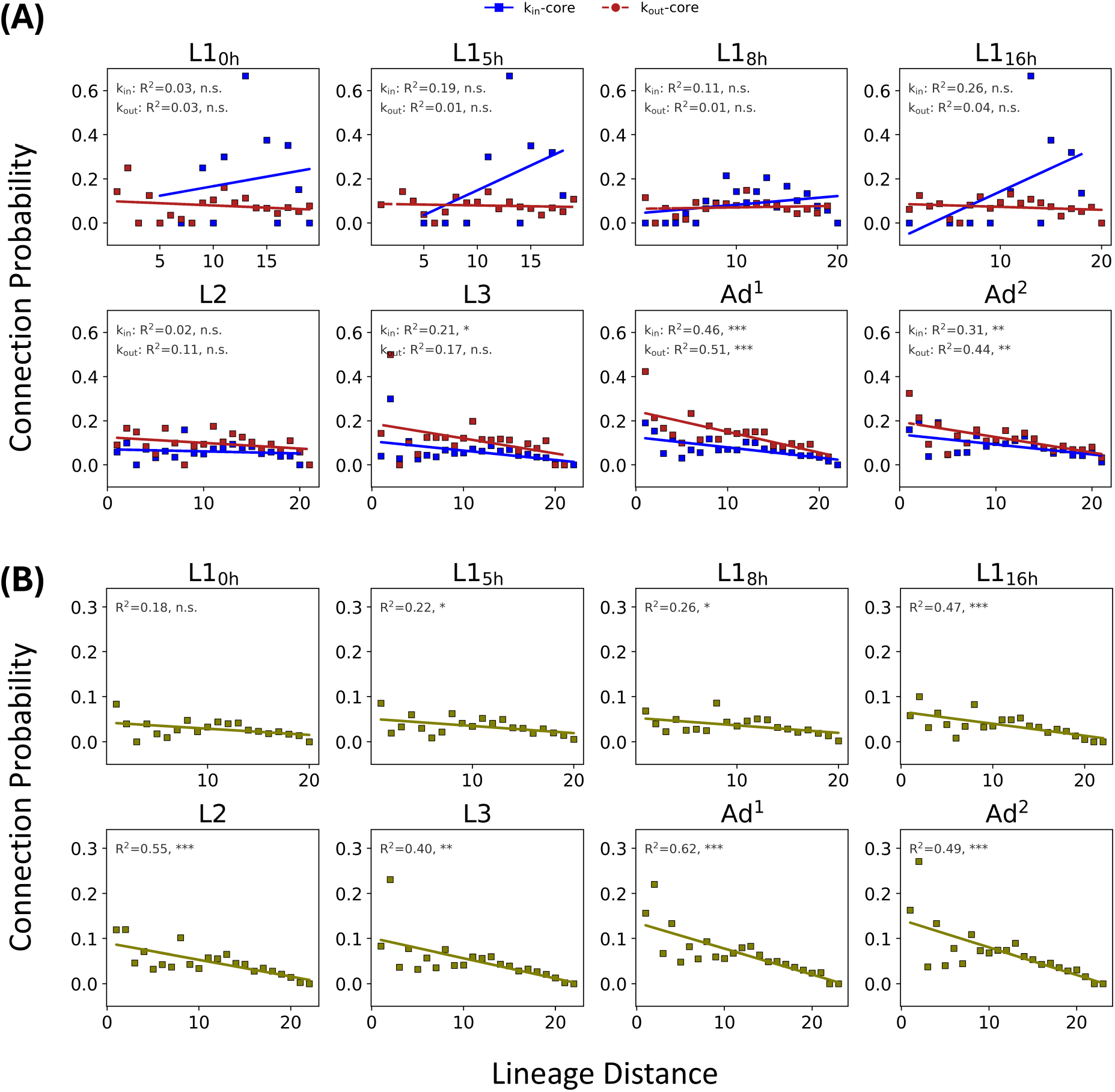
Lineage proximity analysis of the cores and the entire networks. The variation of connection probability with respect to lineage distance is plotted for k-cores (A) and entire networks (B), and linear curve fitting is performed.

We also examined the role of lineage from a different perspective by testing whether core neurons are more closely related to one another than peripheral neurons are among themselves (Figure 4). To answer this, we compute the lineage distance distributions for core and periphery neurons on the four different core decomposition methods for brain networks throughout development. Excluding instances where sample sizes preclude meaningful statistical comparison (especially in the case of out-degree s-core decomposition), we apply the Mann–Whitney U test to evaluate differences between these distributions. We find that in-degree s-cores are much more closely related as attested by moderate values of rank biserial correlation (ranging between 0.44-0.55). In the case of k-core, out-degree-based cores are closely related in the middle stages: from L1-5 to L2, whereas in-degree k-cores are not closely related in any stage. The out-degree s-core is too sparsely populated for comparison in five stages.

**Figure 4:**
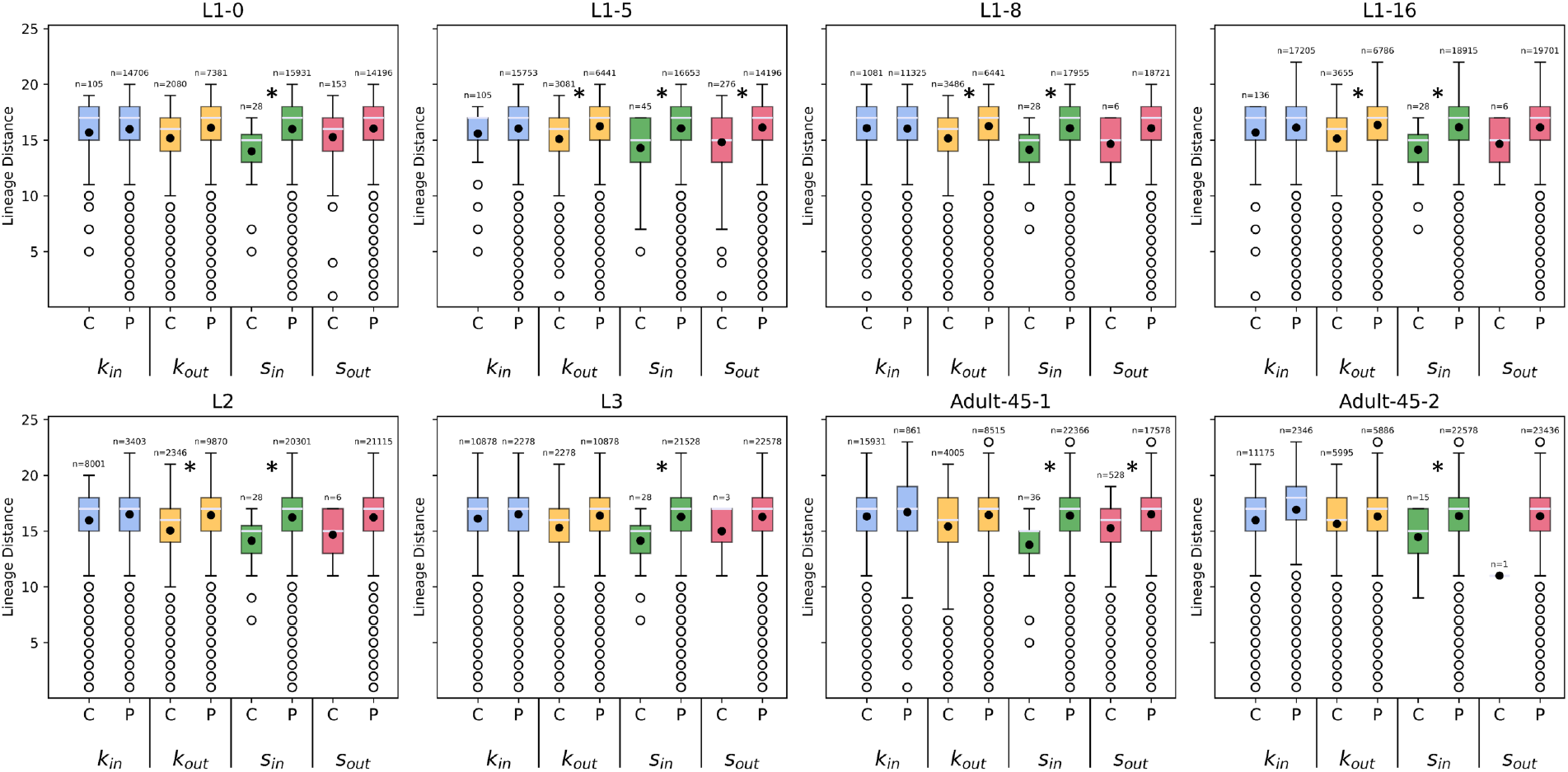
Lineage distance distribution data for core neurons and the peripheral neurons using different core measures across the developmental stages. The star indicates that the distribution for the core neurons is significantly less (Mann-Whitney U test).

Thus, we observe degree-type dependence in the behavior of cores, and these differences are not consistent between k-cores and s-cores, since in s-cores, the in-degree-based s-core is composed of closely related neurons across all stages, while in k-cores, the out-degree-based k-core shows lineage-based proximity, albeit only in the middle stages. The results point to, at most, a partial influence of lineage proximity, further underscoring the fact that multiple factors play a role in synaptic connectivity leading up to the core formation.

#### 2.2.4 Core Topology becomes more Centralized with development

After studying the feature-wise composition, establishing that the various types of core decomposition are more than sufficient to provide a mesoscopic lens, we now probe how the network is organized around the core. To do this, we extend the implementation of Laishram and Soundarajan to directed networks [56]. They created a metric called core centralized score (CE) to measure how centralized the core of a network is. This is implemented on the backbone of the network, that is, the skeletal core subgraph (SCS), which is not unique for a graph. It involves measuring the extent to which the non-core nodes connect with the core nodes. The more their neighbors lie in higher shells, and the higher those shells are, the greater the core centralized score. In the case of a directed network, the connectivity of the non-main core nodes with the core can be probed in two ways (Figure 5). Thus, we computed the CE scores for k-cores for all four possible core-neighbor combinations using an ensemble of 5000 SCSs for each core type. When comparing the two core types, in general, the core topology becomes more centralized with development (Figure 5). However, although CE for in-degree considering both types of edges starts low and ends high with development, the CE for out-degree differs in terms of the edges of non-main core nodes considered. When their incoming edges are considered, the CE is moderate in the early post-embryonic phase and ends up even higher by adulthood, but on the other hand, if outgoing edges are considered, then the CE stays low and flat throughout development. Perhaps this centralized topology is enabled by an increasingly hierarchical organization, which is known to be a key feature of brain networks across taxa, structural [34, 57, 58] and functional both [59–62]. However, it remains to be known how the hierarchical organization changes, if at all, across development and what that means for information flow.

**Figure 5:**
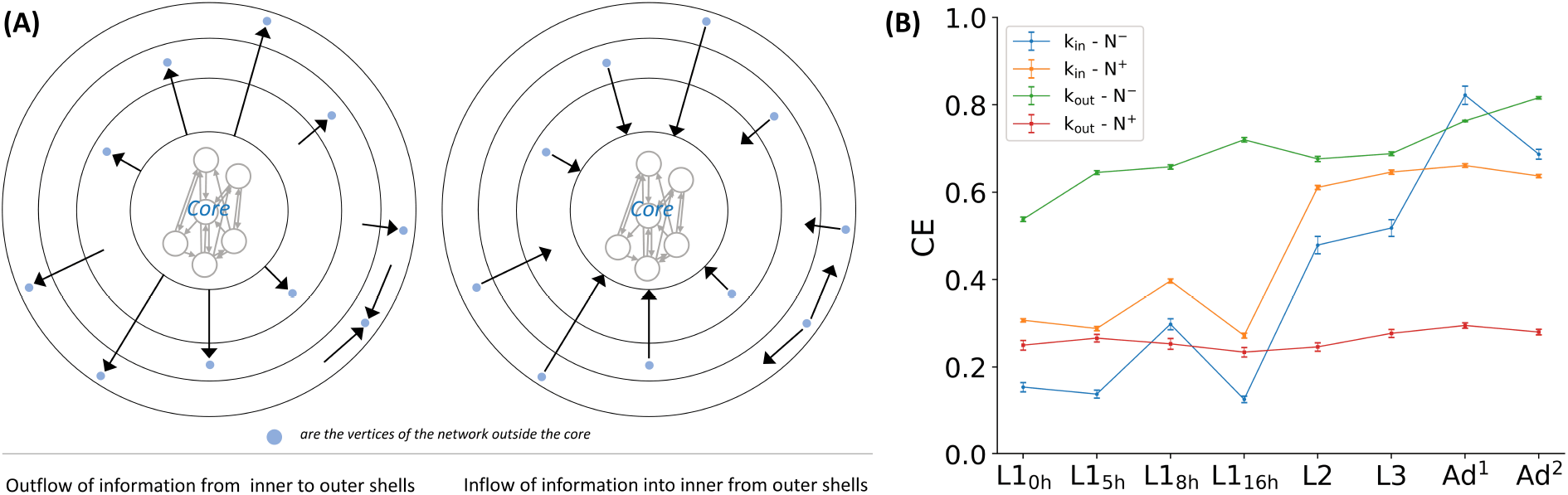
Core centralized scores (CE) of the brain network across development. (A) displays a schematic explaining the directed form of the metric, CE. (B) shows a line plot depicting how core topologies vary across development in terms of their centralization levels. *N*^−^ and *N* ^+^ denote the in-neighbors and out-neighbors, respectively, of nodes not belonging to the main core.

### 2.3 Early and Sustained Presence of Rich Club Organization

To accompany the core analysis, we investigate the presence of rich clubs to holistically study the mesoscopic organization of the connectomes. Like in the core analysis, we studied both weighted and unweighted connectomes, taking directionality of the edges into consideration. As evidenced by the visibly broad regimes where *ϕ*_*norm*_ is significantly greater than 1, the brain connectome consistently exhibits a rich club organization across development for all four degree metrics (Figure 6). Thus, it’s not a feature that emerges at maturation.

**Figure 6:**
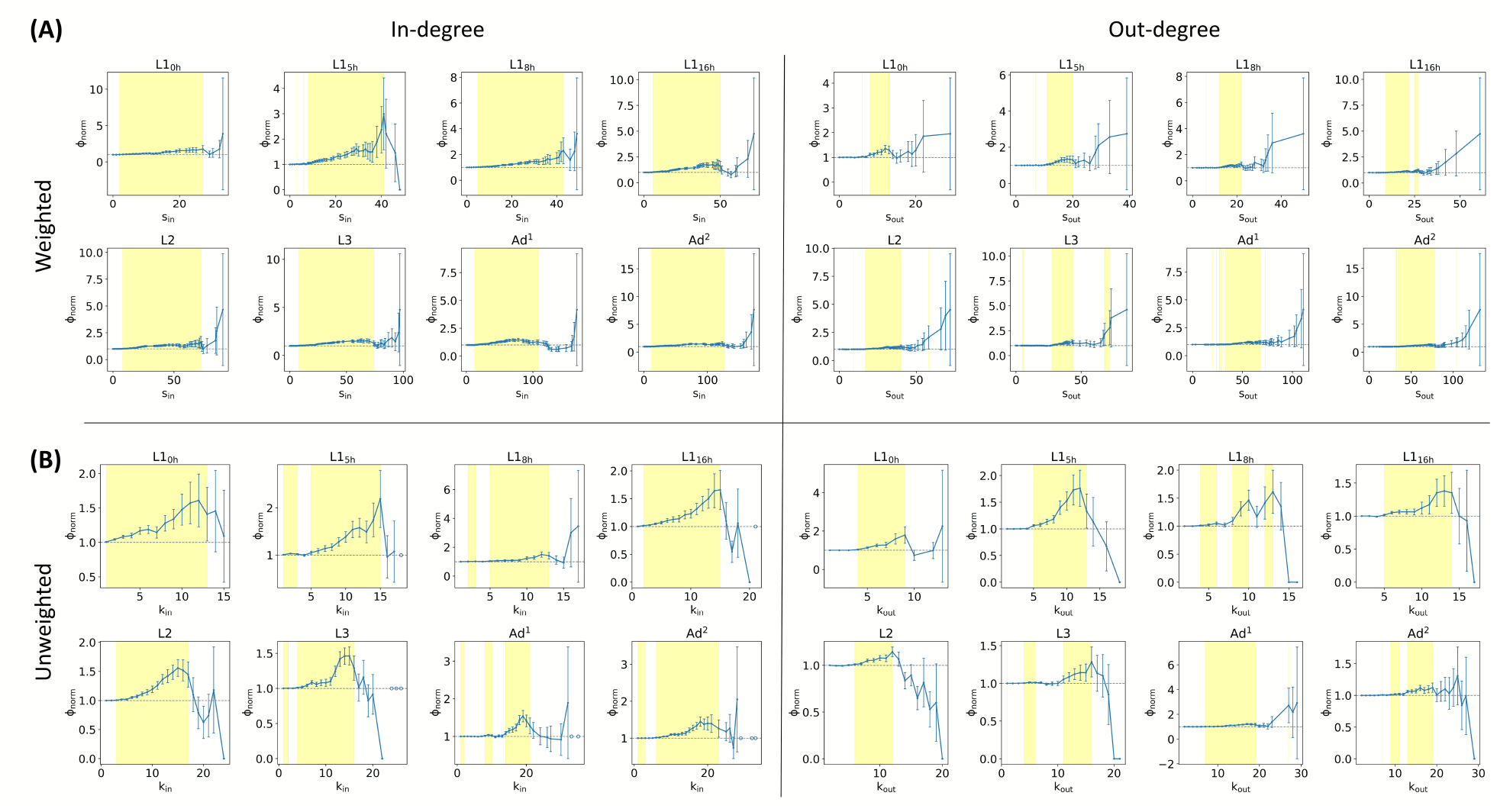
Rich Club Organization. Normalized rich club coefficient is plotted against degree sequences for weighted networks (A) and unweighted networks (B) across development separately for in-degree (left panel) and out-degree (right panel). Yellow spans delineate the rich club regimes, i.e., the range of degrees for which the network shows a significant rich club effect (*ϕ*_*norm*_ *>*= 1+1*×σ*). In cases where the normalized RCC was undefined due to a zero mean *ϕ*_*rand*_, we assign the normalized RCC a value of 1, following Smilkov and Kocarev [63].

The connectomes, however, display heterogeneity across degree-based metrics in their rich club organization (Figure 6). When the weighted in-degree is considered, the rich club regime is broader and also begins earlier in all stages (barring L3). This indicates that nodes receiving strong inputs tend to be more mutually interconnected than those sending strong outputs, even at lower levels of input strength. This suggests that the network structurally favors the formation of tightly coupled input hubs, likely reflecting centralized integration of information from diverse sources. The observations are similar when synaptic strength is disregarded. There is an apparent earlier and longer in-degree rich club regime, meaning that the network supports a deeper or broader nesting of input-receiving subnetworks, where even nodes with just a small number of incoming connections are part of an interlinked structure that gradually intensifies as degree increases. These results suggest that the *C. elegans* connectome contains a broad and gradual hierarchy of input integration where neurons that receive signals, even with modest levels of input, in terms of number or strength of incoming connections, tend to participate in an interconnected backbone of receivers.

To verify the apparent degree-based differences and developmental trends observed in the normalized rich club coefficient plots, we quantify the rich club effect using the area under the curve of the normalized RCC plots (Figure 8). The AUC trend shows that the strength of the weighted rich club effect increases with development, while its unweighted counterpart maintains its level throughout. Thus, the increasing strength of the rich club organization with the development is attributed to the allocation of synapses rather than a remodeling of the connections themselves. Our findings corroborate earlier observations of the human structural brain connectome [64]. Interestingly, the rich club strength when weighted in-degree is considered to be greater than weighted out-degree throughout development, meaning that the cells with strong cumulative input are densely connected with each other at birth, and this connectivity further strengthens as development progresses.

The cores and the rich clubs agree with each other in terms of cell composition (Figure 7). Across the stages, any single core shows a significant overlap with the rich club cohorts, as indicated by the near absence of the unique subsets among the cores in the UpSet plots. Rich clubs show a noticeable presence of unique subsets because of their large size, which is due to the selection of a lower bound of rich club regimes implemented as per consensus in the literature.

**Figure 7:**
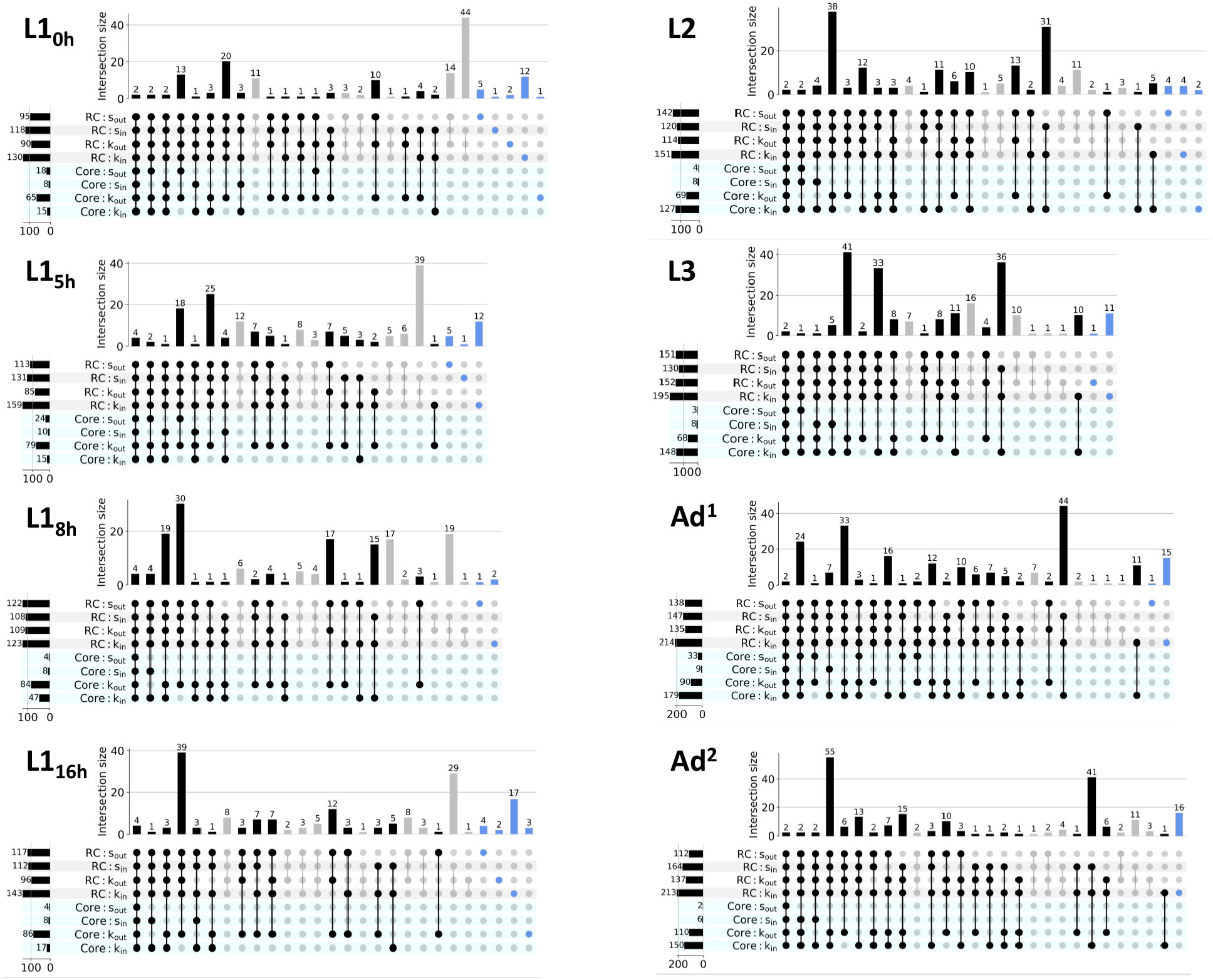
The overlap between the cores and rich clubs for every developmental stage using UpSet plots. Intersections between one or more rich clubs and one or more cores are shown in black connected dots and black bars in the upper panel. Intersections within rich club sets and intersections within core sets are greyed out to emphasize the overlap between core(s) and rich club(s). Moreover, the subsets belonging uniquely to any single rich club or any single core are shown in blue.

**Figure 8:**
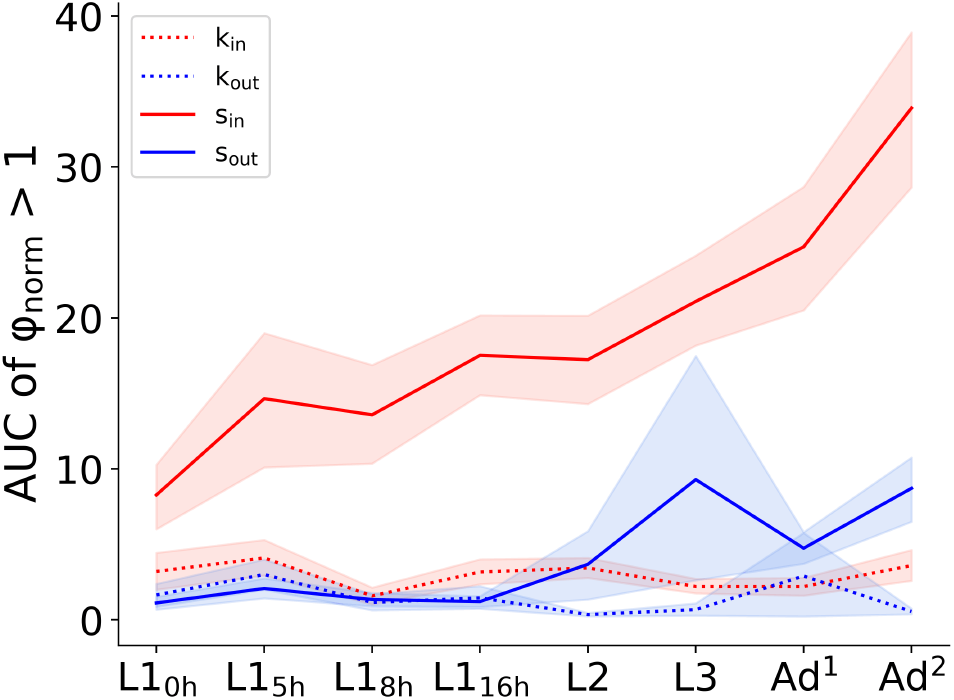
Strength of Rich Club Effect. The line plots depict the strength of the rich club effect in terms of area under the curve for each type of rich club across development. The shaded area corresponding to each line plot shows the 95% confidence interval obtained after Monte Carlo error propagation.

### 2.4 Cohort of Input Hubs is Preserved across Development

The asymmetric synaptogenesis made us contemplate how the cohort of input hubs changes with development. To investigate this, we looked at how the composition of the set of hubs changes with development (Figure 9 (A)). We measure the average consecutive pairwise Jaccard similarities of such sets across the developmental stages. To incorporate the two adult datasets, we take the average of the Jaccard similarity of the L3 stage with the two respective adult datasets. This is done for different percent thresholds of the top degree nodes (5%-50%). Subsequently, we plot the mean consecutive pairwise Jaccard similarity for different thresholds with 95% confidence interval using bootstrapping. It shows that the cohort of input hubs is preserved in comparison to the cohort of output hubs, whether edge strength is accounted for or not.

**Figure 9:**
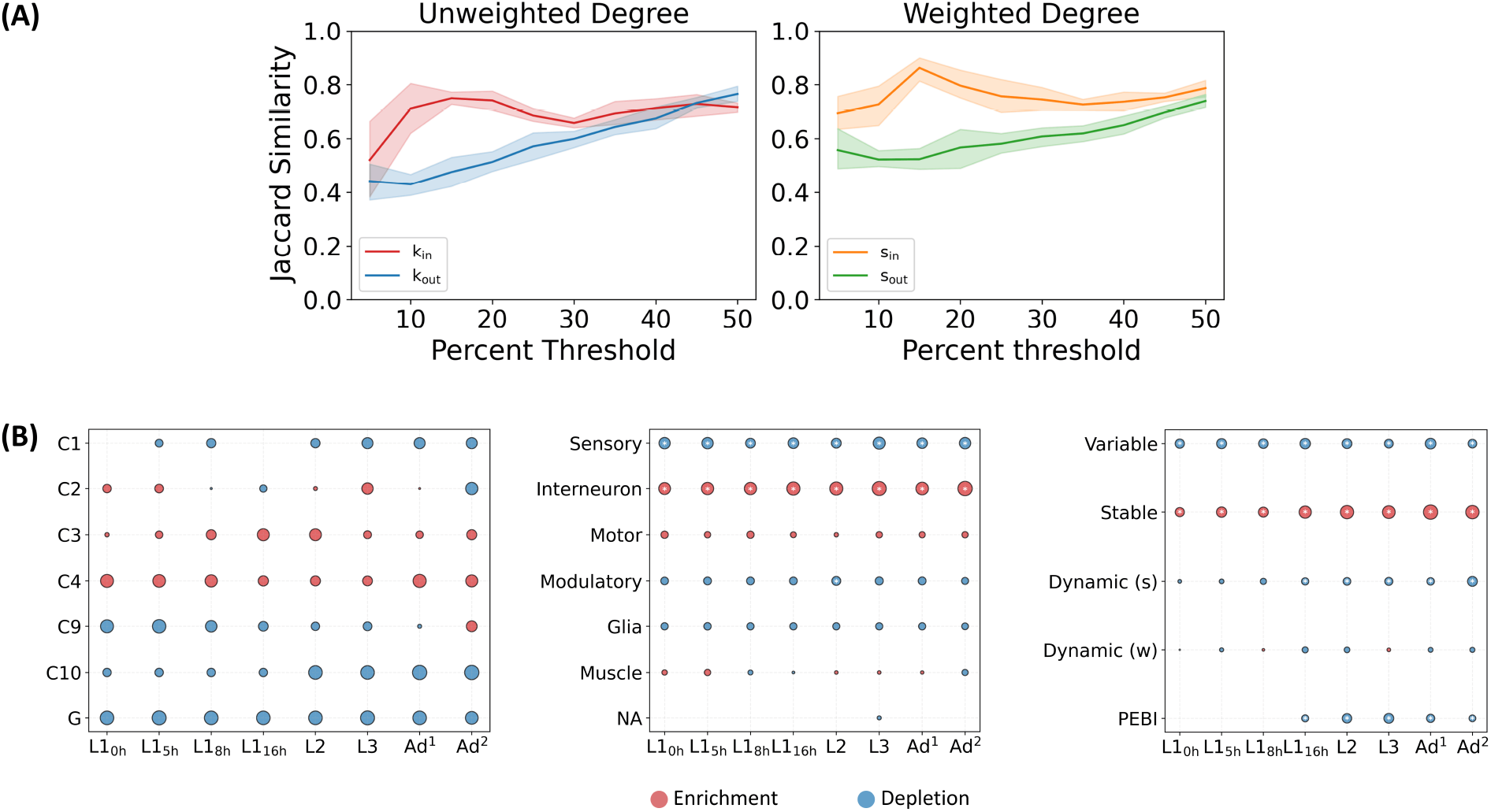
Cohort of Input and Output Hubs. (A) plots the variation of mean pairwise Jaccard similarity between hub sets of consecutive stage-pairs (L3 stage’s Jaccard similarity is taken with respect to both adult datasets and averaged to calculate the overall mean) when connectivity is not weighted (left panel) and when it is weighted (right panel). The shaded areas correspond to 95% confidence interval obtained via bootstrapping. (B) displays bubble charts showing the enrichment of different features in the cohort of top quartile weighted input hubs: community, celltype, edge type, from left to right panels.

The asymmetric synaptogenesis, mesoscopic analysis of the network (the greater overall extent of centralization of in-degree k-core organization and the greater strength of the weighted in-degree rich club effect), and the preservation of a cohort of input hubs across development suggest that the integrative core/cohort is a primary organizing feature of the developing connectome. This prompted us to study the nature of this integrative cohort from a biological standpoint. To do this, we take the top quartile of nodes in terms of their in-degree (ties broken by Katz Centrality), and we study their cell type composition and the behavioral roles they may be biased for by studying their community memberships (Hypergeometric test; FDR correction by Benjamini-Hochberg method) (Figure 9 (B)). Input hubs are consistently enriched for interneurons across all stages (as well as being significantly depleted of sensory neurons), matching their known role in information integration. Witvliet et al. categorized the edges into primarily stable, variable, and developmentally dynamic on the basis of their presence/absence and the degree of variation in their synaptic strength across development. Indeed, they also showed that interneuronal circuits in *C. elegans* are dominated by stable connections, but our analysis refines this observation by showing that such stable connections are specifically enriched among the input hubs that form the integrative backbone. On the other hand, no community showed statistically significant enrichment or depletion among input hubs across development. This indicates that hub membership is broadly distributed across communities, without strong modular bias. The integrative cohort is system-wide, spanning multiple communities rather than confined to a particular behavioral circuit.

## 3 Discussion

We study the growing brain of the nematode, *C. elegans*, comparing and contrasting it at different stages first, on the basis of network parameters and then, mesoscopic organization. Undergoing a continuous development, the *C. elegans* brain displays a weak network connectivity, enabling information integration while still accommodating a hierarchical structure at the expense of strong connectivity. Its edge density progressively increases as it grows. This contrasts with our study of the fruit fly larval and adult connectomes, where we find that the density decreases with development [65]. This disparity is likely due to diverging development patterns: the worm undergoes continuous development while the fly is a holometabolous insect. Consequently, it experiences a remarkable remodeling of its nervous system such that most of its sensory neurons undergo programmed cell death [66–68]. While in the worm most of the brain neurons are formed by the first phase of neurogenesis, in the fruit fly, the second phase, occurring after the release of ecdysone at the end of the third larval stage, triggers multiple types of remodeling that are subtype specific [68]. Building on this context, it should be noted that the fruit fly larva and adult are behaviorally decoupled in addition to possessing distinct ecologies [69].

These reasons also account for the fact that the worm doesn’t undergo a drastic transformation of the core as the fruit fly does. In the *Drosophila* connectome, the core identified is composed entirely of mushroom body neurons in the larval stage but exclusively antennal lobe neurons in the adult stage, indicating a complete turnover of core membership across developmental stages [65]. In contrast, the *C. elegans* connectome exhibits a markedly different pattern across all core definitions considered. A substantial fraction of core neurons are recurrent, consistently reappearing across stages without being permanently retained, while a smaller proportion remains persistent throughout development. These observations suggest that, unlike the fly, the worm brain network retains a degree of continuity in its core organization across development, with neurons repeatedly participating in the core even if not permanently embedded within it.

In contrast to the fly, the roundworm can be understood to develop under a relatively constant ecological background with its central decision-making subnetwork remaining stable throughout its development [29, 70–72]. Under favorable conditions, it has a continuous developmental program progressing through four larval stages to adulthood [70, 71]. Furthermore, the worm displays two locomotory behaviors, crawling and swimming, in its natural habitat of decaying plant matter. Study has revealed that these behaviors are stereotyped across development [17]. It is possible that a stable core facilitates such behavioral stereotypy.

In addition to a consistent core structure across development, the worm brain displays a robust, rich club organization from early on in its post-embryonic development. Due to selective synapse allocation, the weighted rich club effect even strengthens as the brain matures, unlike in the fruit fly [65]. The presence of rich club organization in both the weighted and unweighted brain connectome indicates that both wiring probability and synaptic investment are concentrated within the hub population. Such an arrangement is known to facilitate global communication as well as integration. Our results differ from those of Alstott et al., who reported that the human brain displays a topological rich club but not a weighted rich club [73]. They attributed this phenomenon to the dual functions of the brain: integration and segregation of processing regions. However, two key considerations arise here. First, there is a difference in the resolution of the two types of brain connectomes. Second, even at the neuronal level, connectivity can be characterized in multiple ways, including—but not limited to—chemical synapses, electrical synapses [74, 75], and extrasynaptic signaling [31]. Therefore, it is possible that the brain maintains a balance between integration and segregation through mechanisms beyond chemical synaptic connectivity alone. Further, it’s prudent to understand that the structure and function relationship has various aspects still unknown to us, as evidenced by the dissimilitudes between the two [59, 76].

Notably, the rich club effect across all four degree-based centrality measures is not observed when the topmost cells are exclusively considered, suggesting that each such cell’s connectivity fans outward from the group. Perhaps such an organization serves as a contingency against targeted failures, as certain biological processes operate with a degree of non-randomness. In the case of Alzheimer’s disease, for example, it is understood that, due to their high metabolic activity, hub neurons are predisposed to amyloid-β deposition, which—together with the accumulation of tau filaments within the neurons—constitutes the definitive pathophysiology of the disease [77, 78].

The relatively understudied heterogeneity in terms of degree-based centrality measures in the context of hub-hub connectivity emerges as a potentially important aspect of the system, with implications for future investigation. Given the consistently greater strength of the weighted rich club effect for input hubs, we hypothesize that these neurons may exhibit stronger transcriptional coordination than output hubs [55, 79]. The increasing rich club effect among input hubs with development means that the sum total of synaptic connectivity among integrators increases more than among broadcasters, perhaps in order to facilitate higher-order processing. Given our current understanding of how rich-club organization contributes to neural dynamics, it would be interesting to examine how this picture is refined when directed edges are taken into account [80, 81]. Further, this difference in the meso-scale organization of input and output hubs could be pointing to underlying network features related to the different roles of sources and sinks. Segregation of hubs has been reported in both structural and functional brain networks, providing insights into the overall information flow of the network [82, 83]. The preservation of the cohort of input hubs could possibly be connected to the finding that neuronal activity is correlated with the in-degree [84]. It’s also prudent to take into account that interneurons prominently feature in motifs containing clustered synapses, which are involved in compartmentalized computations [85]. Perhaps, a more extensive rewiring of the cohort would thus perturb the robustness of the brain network’s function. Interestingly, even in the Dauer connectome, among all the cell types, the connectivity of interneurons is preserved the most [30]. On the macroscale level, we believe our results on the directional asymmetry of the core topology in terms of its centralization could potentially be useful in the study of metastable brain waves of directed connectomes, since it is already known that degree hubs tend to act as sources of brain waves [86].

Witvliet et al. reported a biased process of synaptogenesis in favor of the input hubs [29]. We show that this asymmetry extends onto higher scales in that the input hubs seem to be a primary organizing feature around which the brain network develops. The increasingly cohesive weighted in-degree rich club facilitates a meso-scale consolidation of the integrative core/cohort. Further, we find that the cohort of input hubs is preserved, enriched in interneurons, and consequently, stable connections. Finally, the indegree core organization becomes progressively more centralized, giving rise to a macro-scale establishment of a hierarchical network topology.

## 4 Methods

### Data Collection

The synaptic connectivity data for 8 different brain connectomes of *C. elegans* across 7 unique timepoints in its development are taken from Witvliet et al. [29]. Accompanying this are the celltype information and the classification of the connections. To contextualise the appearance of transient and recurrent core cells, we use birth time data of neurons (except CANR) from [36] downloadable from their lab webpage. The birth time data for the remaining cells (CANR, glia and muscles) is obtained from wormweb.org. All birth time data originially comes from Sulston et al. 1977 and 1983 [5, 87]. The times are measured from the fertilization which occurs 50 minutes prior to the first cleavage [87]. For the lineage study, we compile the data from [88] (for the neurons) and [89] (for the glia and muscles) both of which ultimately source their data from the aforementioned pioneering work by Sulston and his group. The community dataset is taken from Emmons’ 2024 work [37]; it was generated by applying spectral method of Leicht and Newman [90] onto connectivity matrix weighted by synapse size of chemical synapses and gap junctions.

### Network Properties

Each brain network can be represented as an adjacency matrix where *A*_*ij*_ gives the strength of synaptic connection (number of synapses) going from cell i to cell j. In the case of unweighted analyses, the binarized matrix was considered meaning any non-zero *A*_*ij*_ value was taken to be 1. In the early stages, there are isolated nodes representing the cells that begin their synapses formation during the post-embryonic phase. The network density, which measures the degree of connectivity among the nodes, is given by 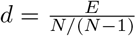 where *E* is the number of connections and *N* is the number of nodes in the network. Weak and strong connectivity are calculated after removing the isolated nodes. The degree is the number of incoming and outgoing connections incident on node *i*. It is given by 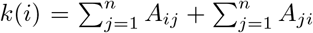 self-loops (11) exist only in *Ad*^1^ stage and are excluded from analyses to ensure methodological consistency and fair cross-stage comparison. Lastly, the average connection weights is given by 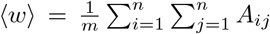. All structural properties are calculated under the framework of networkx library [91].

### k-core decomposition

A k-core decomposition is a process that iteratively simplifies a graph by examining node degrees. For a given graph *G*, the k-core is defined as the maximal subgraph in which every vertex has degree at least k. To obtain it, vertices with degree smaller than k are repeatedly removed, since their deletion may reduce the degrees of neighboring vertices and trigger further removals. As k increases, this pruning procedure produces a hierarchy of progressively smaller, nested subgraphs, each corresponding to a different k-core.

### s-core decomposition

The s-core generalizes the k-core concept to weighted networks [92]. For a weighted graph *G*, the s-core is defined as the maximal subgraph in which each node has a weighted degree (strength) of at least s. The decomposition is obtained through an iterative pruning process: nodes whose weighted degree falls below s are repeatedly removed, as their removal can reduce the strengths of adjacent nodes and lead to further eliminations. By increasing s, this procedure yields a sequence of nested subgraphs, known as s-cores.

### Core Centralized Score of the Core

For each of the *k*_*in*_ and *k*_*out*_ core topologies, we create an ensemble of 5000 skeletal core sub-graphs (SCSs), then using the formula (modified to account for directionality) [56] 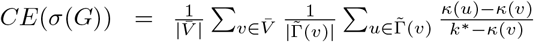 take an average of the core centralized score score across ensemble for each stage. In the equation, *σ*(*G*) refers to a skeletal core subgraph of the entire graph, 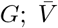 is the set of nodes outside the main core; 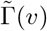 is the set of neighbors of *v* in the core subgraph of *v* (denoted by *κ*(*v, G*) *core*); *κ* gives the core number of a node; and *k*^*\**^ is the degeneracy of the entire graph, *G* i.e. the main core number. To incorporate directionality, we calculate *CE* by separately assessing the predecessors 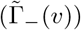 and successors of 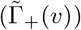 node v for a given core type.

### Community Detection

To robustly obtain the communities we run 100 iterations of the Louvain algorithm using different random seed values each time at the default resolution. Then, we create a co-occurrence matrix quantifying the number of times any two nodes feature in the same community. Finally, we run another instance of Louvain algorithm on this matrix to get the final list of communities. The modularity gain is calculated from the following formula [93]: 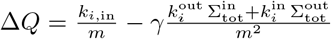where 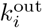 and 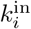 are the outer and inner weighted degrees of node *i*, and 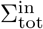 and 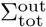 represent the sum of the incoming and outgoing edges incident to nodes in *C*.

### Rich Club Analysis

Rich club measures the extent to which high degree nodes are connected with each when accounting for random chance. By taking direction into consideration, we carry out rich club analysis on both unweighted (binarized) and weighted brain networks. To normalize the so obtained rich club coefficient we employ degree preserved and weighted-degree (strength) preserved randomized null models. For directed unweighted rich club coefficient, we use the formula:

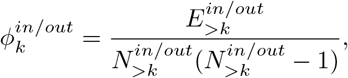

where, 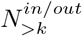 is the number of nodes with in/out-degree greater than *k* and 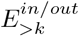 is the number of directed edges between them. In the case of weighted rich club evaluation, we quantify the weighted connectedness (the sum of the weights of the edges between the ‘rich’ nodes).

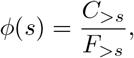

where, *C*_*>s*_ is the weighted connectedness of nodes with weighted degree greater than *s* and *F*_*>s*_ is their maximum possible weighted connectedness. Using appropriate null models renders the exact formulation of *F*_*>s*_ unnecessary since it will be the same for the empirical and null model due to the sequence of nodes according to their weighted degree being the same [73]. Thereby, the normalized weighted rich club coefficient is given by 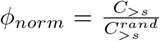 where 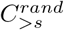 is the weighted connectedness of a weighted-degree-preserving null model. This framework is then simply applied separately onto weighted in-degree and weighted out-degree sequences of a directed and weighted network.

For each degree threshold k, the RCC was computed from the subgraph induced by nodes with weighted degree strictly greater than k, consistent with methods described in the literature [44, 73, 94, 95]. These regimes—whether contiguous or disjoint—yield multiple thresholds with significant rich-club organization. Since increasing thresholds select increasingly exclusive sets of high-degree nodes, the corresponding rich-club subgraphs are naturally nested [80]. Sometimes, we obtain not a regime per se but a single, isolated threshold in the degree sequence—or a ‘singleton’.

We identify rich-club regimes as the ranges of degrees where the *ϕ*_*norm*_ *>*= 1 + 1 *× σ*, with *σ* representing the standard deviation of the normalized RCC. This follows the approach used in Towlson et al. (2013) [44]. Also in line with same work, we define the nodes forming a rich club using the lowest degree threshold for the first regime (or singleton) of a network thereby, capturing the largest associated rich-club subgraph.

### Quantification of the Rich Club effect

To quantify the rich club effect, we calculate the area under the curve (AUC) of the normalized RCC plot of a network above the *ϕ*_*norm*_ = 1 line using the trapezoid rule. In the case of singletons, we carry out an approximation by taking the magnitude of by which *ϕ*_*norm*_ is greater than 1 for that threshold value. The 95% confidence interval was calculated using Monte Carlo Error Propagation method (1000 iterations).

### Randomized Null Models

To normalize the rich club coefficients we create an ensemble of degree-preserved randomized null modesl as well as weighted-degree-preserved (strength) randomized null models. For both, we use the simulated annealing method using the code provided by Milisav et al. [96].

### Cohort of Hubs

We decide upon using top quartile of nodes in terms of their in-degree (ties broken by katz centrality) as the representative cohort after rejecting alternative statistical methods for identifying in-degree hubs— z-score thresholding: rejected because degree distributions aren’t Gaussian; power-law tail cutoff: rejected because the networks can’t be proven to be fat-tailed; and the rank-degree curve method: rejected for being unstable and inconsistent across stages. Thus, we settle on a fixed top-percentile rule for consistency.

### Packages Used for Analysis

Here is the exhaustive list of all the Python packages we used to carry out this study: networkx [91], numpy [97], pandas [98], and scipy [99] for the analyses and matplotlib [100], and seaborn [101] for the visualizations.

## 5 Acknowledgement

A.S. acknowledges the Department of Science and Technology (DST), Government of India, for financial support through the DST INSPIRE Faculty Grant No. IFA-21-PH-276. P.Y. acknowledges the University Grants Commission (UGC), Government of India, for financial support under the Junior Research Fellowship scheme. The authors thank Aiwin T. Vadakkan for his assistance during the initial stages of the project, and the lab members for their timely help and valuable discussions. The authors also gratefully acknowledge IISER Tirupati for providing access to the High-Performance Computing facility.

## Notes

### Competing Interest Statement

The authors have declared no competing interest.

### Summary of Updates

This revised version includes minor textual updates to the Discussion section to provide additional context.

